# The USTC complex co-opts an ancient machinery to drive piRNA transcription in *C. elegans*

**DOI:** 10.1101/377390

**Authors:** Chenchun Weng, Asia Kosalka, Ahmet C. Berkyurek, Przemyslaw Stempor, Xuezhu Feng, Hui Mao, Chenming Zeng, Wen-Jun Li, Yong-Hong Yan, Meng-Qiu Dong, Cecilia Zuliani, Orsolya Barabas, Julie Ahringer, Shouhong Guang, Eric A. Miska

**Affiliations:** Hefei National Laboratory for Physical Sciences at the Microscale, School of Life Sciences, University of Science and Technology of China, Hefei, Anhui 230027, P.R. China.; Wellcome Cancer Research UK Gurdon Institute, University of Cambridge, Tennis Court Road, Cambridge CB2 1QN, UK; Department of Genetics, University of Cambridge, Downing Street, Cambridge CB2 3EH, UK.; National Institute of Biological Sciences, Beijing 102206, China.; Wellcome Sanger Institute, Wellcome Genome Campus, Cambridge CB10 1SA, UK; Structural and Computational Biology Unit, European Molecular Biology Laboratory (EMBL), 69117 Heidelberg, Germany.

**Keywords:** piRNA, Ruby motif, SNPC, PRDE-1, SNPC-4, TOFU-4, TOFU-5, snRNA, U6 RNA

## Abstract

Piwi-interacting RNAs (piRNAs) engage Piwi proteins to suppress transposons and non-self nucleic acids, maintain genome integrity, and are essential for fertility in a variety of organisms. In *C. elegans* most piRNA precursors are transcribed from two genomic clusters that contain thousands of individual piRNA transcription units. While a few genes have been shown to be required for piRNA biogenesis the mechanism of piRNA transcription remains elusive. Here we used functional proteomics approaches to identify an upstream sequence transcription complex (USTC) that is essential for piRNA biogenesis. The USTC complex contains PRDE-1, TOFU-4, TOFU-5 and SNPC-4. The USTC complex form a unique piRNA foci in germline nuclei and coat the piRNA cluster genomic loci. USTC factors associate with the Ruby motif just upstream of type I piRNA genes. USTC factors are also mutually dependent for binding to the piRNA clusters and to form the piRNA foci. Interestingly, USTC components bind differentially to piRNAs in the clusters and other non-coding RNA genes. These results reveal USTC as a striking example of the repurposing of a general transcription factor complex to aid in genome defence against transposons.

## INTRODUCTION

Piwi-interacting RNAs (piRNAs) are a class of small (21-30 nucleotide) RNAs that associate with Piwi proteins, a highly conserved subclass of Argonaute proteins, and play significant roles in fertility and genome stability (Carmell et al. 2007; Cox et al. 1998; Houwing et al. 2007; Lin and Spradling 1997; Palakodeti et al. 2008; Das et al. 2008; Girard et al. 2006; Batista et al. 2008). piRNAs program genome elimination during the sexual reproduction in ciliates (Chalker and Yao 2011) and engage in sex determination, virus defense, regeneration and neoblast function in animals (Palakodeti et al. 2008; Kiuchi et al. 2014; Reddien et al. 2005; Schnettler et al. 2013; Zhao et al. 2013). piRNAs also play important roles in male fertility in humans (Gou et al. 2017). In *C. elegans*, piRNAs preserve genome integrity in the germ line by recognizing and silencing “non-self” genomic loci such as transposons or other foreign nucleic acids and induce chromatin modifications (Luteijn and Ketting 2013; X. Feng and Guang 2013; Malone and Hannon 2009; Mao et al. 2015; Ashe et al. 2012; Bagijn et al. 2012). In addition to the function of piRNAs in the germline, a recent report provided evidence for the presence of piRNA factors such as PRDE-1 and PRG-1 in *C. elegans* neuronal cells (Kim et al. 2018).

*C. elegans* piRNAs are also termed as 21U-RNAs, given that they are predominantly 21 nucleotides (nt) in length and have a strong bias towards 5’-monophosphorylated uracil *(Das et al. 2008; Batista et al. 2008; Bagijn et al. 2012; Gu et al. 2012)*. 21U-RNAs silence transposons and protein-coding genes independently of the endonuclease or slicing activities of the Piwi protein PRG-1. Instead, 21U-RNAs scan for foreign sequences, while allowing mismatched pairing with the targeted mRNAs (Ashe et al. 2012; Bagijn et al. 2012; Shirayama et al. 2012). Upon targeting, the 21U-RNA/PRG-1 complex recruits RNA-dependent RNA polymerase (RdRP) to elicit the generation of secondary small interfering RNAs (siRNAs), referred to as “22G-RNAs”. The 22G-RNAs are then loaded onto worm-specific Argonaute proteins (WAGOs) to conduct gene silencing processes (Mao et al. 2015; Ashe et al. 2012; Bagijn et al. 2012; Shirayama et al. 2012; Lee et al. 2012). Meanwhile, “self” mRNAs are protected from 21U-RNA-induced silencing by the CSR-1 Argonaute pathway (Shirayama et al. 2012; Conine et al. 2013; Seth et al. 2013). Therefore, 21U-RNAs are required to initiate the epigenetic silencing, yet the inheritance of the silencing requires 22G-RNAs (Das et al. 2008; X. Feng and Guang 2013; Bagijn et al. 2012; Batista et al. 2008; Gu et al. 2012; Ruby et al. 2006).

Unlike siRNAs and miRNAs, the biogenesis of piRNAs is a Dicer-independent process (Siomi et al. 2011; Grishok 2013; Klattenhoff and Theurkauf 2008). In *Drosophila*, the long piRNA precursors are transcribed from two different genomic sources, the uni-strand and dual-strand clusters, which are further processed and amplified by distinct factors to conduct their respective functions (Handler et al. 2011; Vourekas et al. 2015; Mohn et al. 2014; Klattenhoff et al. 2009; Brennecke et al. 2007; Goriaux et al. 2014; Pane et al. 2011; Czech and Hannon 2016; Ipsaro et al. 2012; Saito et al. 2007). The mouse piRNAs are classified into pre-pachytene and pachytene piRNAs based on their expression patterns (X. Z. Li et al. 2013; Aravin et al. 2008), pachytene piRNA transcription distinctively requiring the transcription factor A-MYB (X. Z. Li et al. 2013).

21U-RNAs are expressed in the germ line from thousands of genomic loci, and mostly from two large genome clusters on chromosome IV. They are first transcribed by RNA polymerase II that initiate precisely 2 nt upstream of the 5’ end of mature 21U-RNAs to generate 25-29 nt capped small RNA (csRNA) precursors (Gu et al. 2012; Cecere et al. 2012; Weick et al. 2014). Then, the precursors are decapped at the 5’-end, the first two nucleotides are removed, and the extra-nucleotides at 3’-ends are trimmed off and methylated, to produce mature 21U-RNAs (Tang et al. 2016; Montgomery et al. 2012; Billi et al. 2012; de Albuquerque et al. 2014; Weick et al. 2014). C. *elegans* encodes two Piwi proteins, PRG-1 and PRG-2. Whereas the function of PRG-2 is unknown, mature 21U-RNAs associate with PRG-1 to conduct their functions (Das et al. 2008; Bagijn et al. 2012; Batista et al. 2008). The binding of 21U-RNAs to PRG-1 is important for their production and silencing effect of 21U-RNAs. 21U-RNAs are absent in *prg-1* mutant animals (Gu et al. 2012; Wang and Reinke 2008) and loss of 21U-RNA/PRG-1 complexes leads to reduced fertility (Das et al. 2008; Batista et al. 2008; Wang and Reinke 2008). Interestingly, untrimmed piRNAs with 3’ extensions are stable and associate with PRG-1, yet they are unable to robustly recruit other downstream factors, and therefore compromise the silencing effect (Tang et al. 2016).

Two types of piRNAs have been described in *C. elegans*. Type-I 21U-RNAs are predominantly transcribed from two broad regions on chromosome IV and contain an eight nt upstream Ruby motif (CTGTTTCA) and a small YRNT motif in which the T corresponds to the first U of the 21U-RNA. Type-ll 21U-RNAs are present outside of chromosome IV and lack the Ruby motif (Gu et al. 2012; Ruby et al. 2006). Each 21U-RNA is independently transcribed as a short RNA precursor by RNA polymerase II. Recent work identified PRDE-1 and SNPC-4 in a complex that binds to the Ruby motif of type I piRNA loci which are essential for the transcription of piRNA precursors (Weick et al. 2014; Kasper et al. 2014). A Forkhead family transcription factor, *unc-130*, was shown to bind 21U-RNA promoters and has been implicated in piRNA transcription (Cecere et al. 2012). PID-1 may function to promote piRNA processing in the cytoplasm (de Albuquerque et al. 2014). A genome-wide RNAi screen identified seven Twenty-One-u Fouled Ups (TOFUs) that are engaged in distinct expression and processing steps of 21U-RNAs (Goh et al. 2014). However, it is still unclear of how the transcription of 21U-RNA precursors is controlled. Here, we used functional proteomics and identified the upstream transcription complex USTC containing PRDE-1, SNPC-4, TOFU-4, and TOFU-5 which bound to the promoters of 21U-RNA precursors to drive their expression in the *C. elegans* germline.

## RESULTS

### Mass spectrometry and yeast-two-hybrid screens identify a complex containing PRDE-1, SNPC-4, TOFU-4 and TOFU-5 proteins

Our previous study identified *prde-1* (piRNA silencing-defective) as being essential for 21U-RNA generation in *C. elegans (Weick et al. 2014)*. Later, it was found that a small nuclear RNA activating protein complex, SNPC-4, interacts with PRDE-1 and promotes piRNA generation (Kasper et al. 2014; Goh et al. 2014). To further understand the transcription regulation of piRNAs, we searched for proteins that interact with PRDE-1 and SNPC-4 using co-immunoprecipitation mass spectrometry (co-IP-MS) and yeast-two-hybrid (Y2H) methods. The top candidates interacting with PRDE-1 and SNPC-4 are listed in Supplementary Table S1. In line with previous results, SNPC-4 was shown to interact with PRDE-1 by co-IP-MS (Supplementary Table S1). Interestingly, we found that TOFU-4, a protein that is known to be required for piRNA biogenesis from the previous genome-wide RNAi screening, interacts with PRDE-1 by IP-MS and Y2H. The Y2H experiment of SNPC-4 only isolated TOFU-5 (Supplementary Table S1).

To confirm these protein-protein interactions using independent means, we generated single-copy GFP-3xFLAG tagged TOFU-4, and TOFU-5 (abbreviated as TOFU-4::GFP and TOFU-5::GFP respectively) transgenic strains using the Mos1-mediated single-copy insertion (MosSCI) technology (Frøkjaer-Jensen et al. 2008). The TOFU-4::GFP and TOFU-5∷GFP transgenes rescued the *tofu-4* and *tofu-5* mutant phenotypes, respectively, suggesting that the tagged proteins could recapitulate the functions of endogenous proteins (Fig. S1A,B). IP-MS of TOFU-5 identified the SNPC proteins, SNPC-1.1, −3.1, −3.2, −3.4, and −4 as the top candidates of TOFU-5 interactors (Table 1). The identification of SNPC-4 in the IP-MS experiment of TOFU-5 is consistent with the Y2H result of SNPC-4 (Table 1). The proteomic experiments among the four proteins are summarized in Fig. 1A.

**Figure 1.**
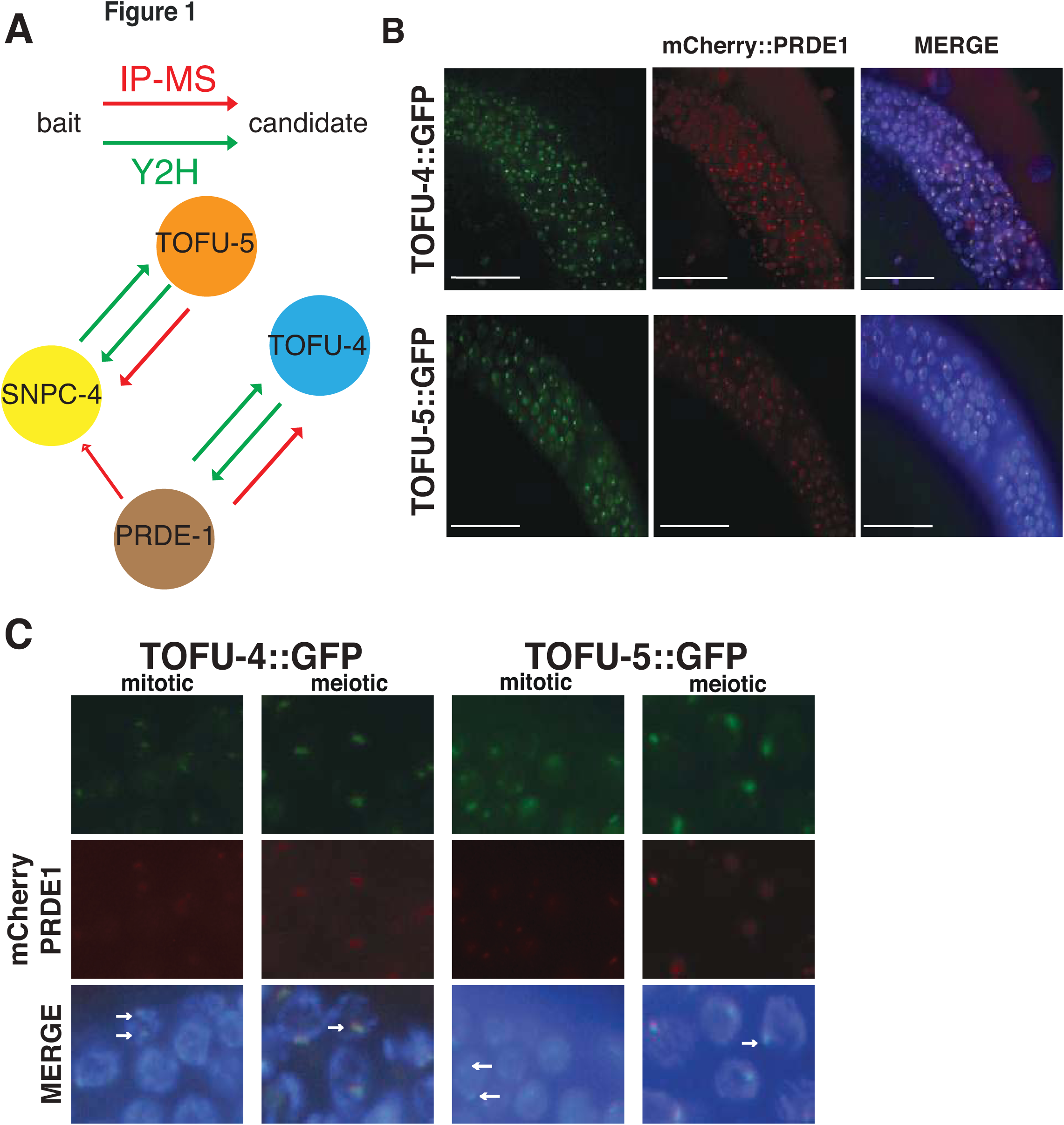
Functional proteomics identify the USTC complex. (A) Summary of protein-protein interaction experiments between PRDE-1, TOFU-4, TOFU-5, and SNPC-4. (B,C) Subcellular colocalization of TOFU-4∷GFP and TOFU-5∷GFP (green) with mCherry:PRDE-1 (red) in young adult germline nuclei (B) and zoomed images from the mitotic zone and the meiotic zone (C).

Both TOFU-4∷GFP and TOFU-5∷GFP exhibit distinct foci in the germline nuclei (Fig. S2A,B). Previous work showed that PRDE-1 colocalizes with SNPC-4 and forms distinct foci in germline cell nuclei (Weick et al. 2014; Kasper et al. 2014). We immunostained PRDE-1 with anti-PRDE-1 antibody in TOFU-5∷GFP animals and found that PRDE-1 and TOFU-5 also colocalize with each other in the germline nuclei (Fig. S2C). To further confirm the protein-protein interactions of PRDE-1, TOFU-4 and TOFU-5 proteins, we crossed the mCherry∷PRDE-1 transgene into TOFU-4∷GFP and TOFU-5∷GFP expressing animals (Fig. 1B,C). Consistently, both TOFU-4 and TOFU-5 are colocalized with PRDE-1 as distinct foci in germ cell nuclei. Importantly, nuclei in the mitotic zone exhibited two foci and nuclei in the meiotic zone one focus consistent with the ploidy of the cells (Kasper et al. 2014) (Fig. 1C). Therefore, we conclude that PRDE-1, SNPC-4, TOFU-4, and TOFU-5 likely function as a protein complex to engage in piRNA biogenesis.

### PRDE-1, SNPC-4, TOFU-4, and TOFU-5 coat piRNA clusters and are enriched at the Ruby motif

SNPC-4 has been previously shown to associate with the chromosome IV piRNA clusters and that binding depends on the presence of PRDE-1 (Kasper et al. 2014; Weick et al. 2014). To test whether PRDE-1, TOFU-4 and TOFU-5 co-localise with SNPC-4 at piRNA clusters, we performed ChIP-seq experiments with all four factors in young adults when piRNA expression is at its peak (Das et al. 2008; Batista et al. 2008); all experiments were done in duplicates (Fig. S3). We found that the four factors have similar genome-wide binding profiles, with strong enrichment on chromosome IV piRNA clusters (Fig. 2A,B, Fig. S4A,B). Curiously, SNPC-4 did not appear enriched on the smaller piRNA cluster (cluster I), however, this might be a reflection of lower signal to noise ratio of the SNPC-4 ChIP-seq experiments (Fig. 2A). Finally, TOFU-4, TOFU-5, SNPC-4, and PRDE-1 all “coated” piRNA genes broadly, showing signal around piRNA genes above the genome average and a peak upstream of the transcription start site (Fig. 2C).

**Figure 2.**
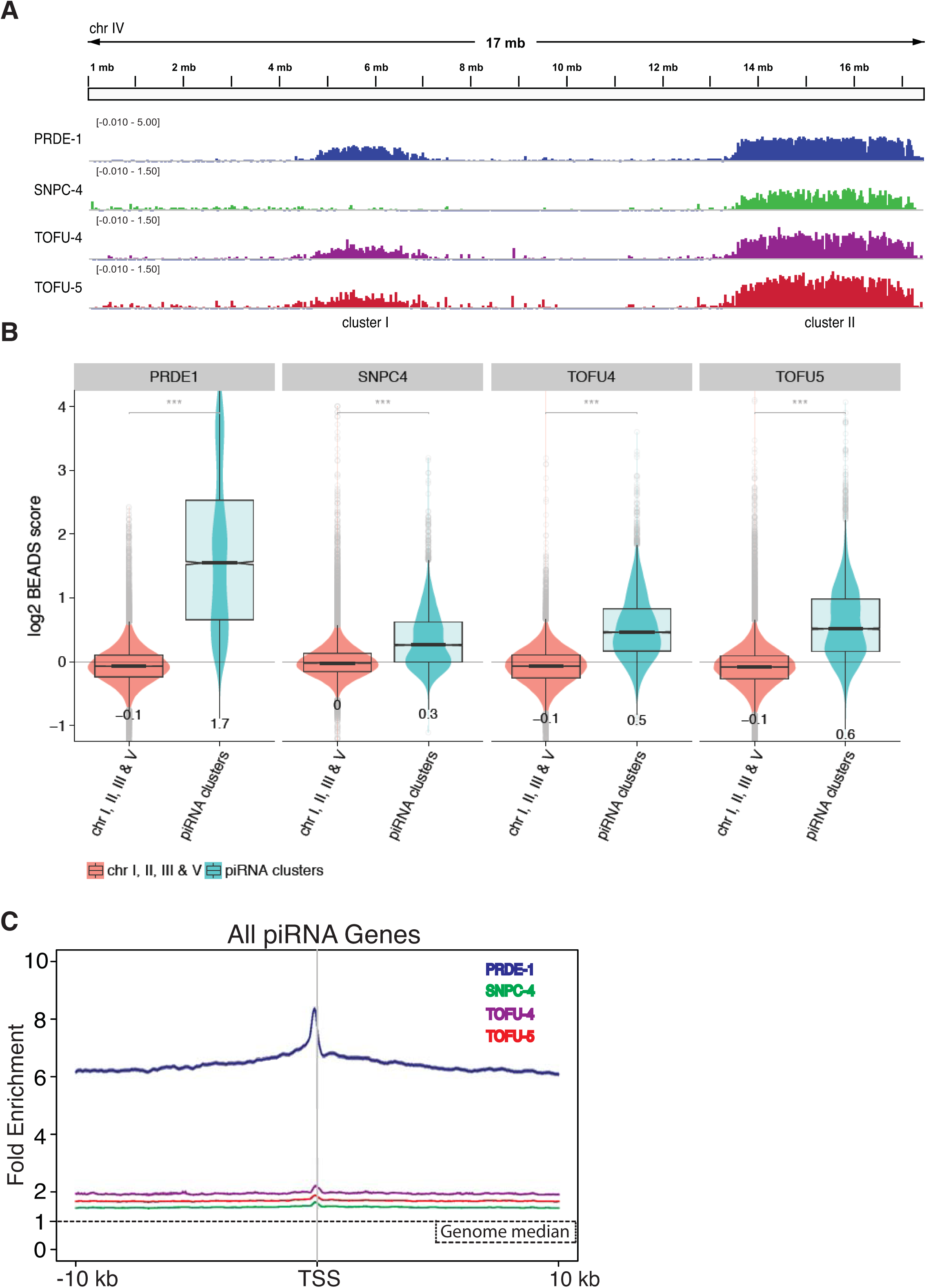
The USTC factors coat the piRNA clusters of chromosome IV. (A) The binding profiles of USTC factors across chromosome IV. ChIP signal was normalized with rBEADS and log2 transformed. (B) Quantification of ChIP-seq signal on chromosome IV piRNA clusters and other somatic chromosomes (l,ll,lll,V). The signal was calculated in 1kb bins. All USTC factors are significantly enriched on piRNA clusters. (C) Enrichment profile of USTC factors around the transcription start site (TSS) of all piRNA genes. The plot is anchored on the 1st U nucleotide of each piRNA.

*C. elegans* piRNAs are categorized into type I and type II piRNAs. Type I piRNAs feature the Ruby motif upstream of the TSS of each piRNA transcription unit are mostly found in the two chromosome IV piRNA clusters whereas type II piRNAs are more distributed and lack the Ruby motif (Gu et al. 2012; Ruby et al. 2006). To ask if TOFU-4, TOFU-5, SNPC-4, and PRDE-1 preferentially bound a class of piRNA gene, we plotted heat map profiles of the USTC components around type I and type II piRNA TSS respectively (Fig. 3A, S5). We observed that all four factors exhibit robust enrichment around type I piRNA genes (Fig. 3A) and that z-scores overlapped the Ruby motif (Fig. 3B). In contrast, PRDE-1 showed little enrichment at type II piRNAs (Fig. S5A,B) consistent with our previous finding that PRDE-1 is not required for type II piRNA transcription (Weick et al. 2014). SNPC-4 and TOFU-5 however were enriched at type II piRNA genes (Fig. S5A,B). Combining the proteomic experiments, subcellular colocalization, and the enrichment at type I piRNA genes, we conclude that TOFU-4, TOFU-5, SNPC-4, and PRDE-1 function as a protein complex to promote the biogenesis of piRNAs by binding to the upstream sequence of piRNA transcription. We therefore named this complex the upstream sequence transcription complex (USTC).

**Figure 3.**
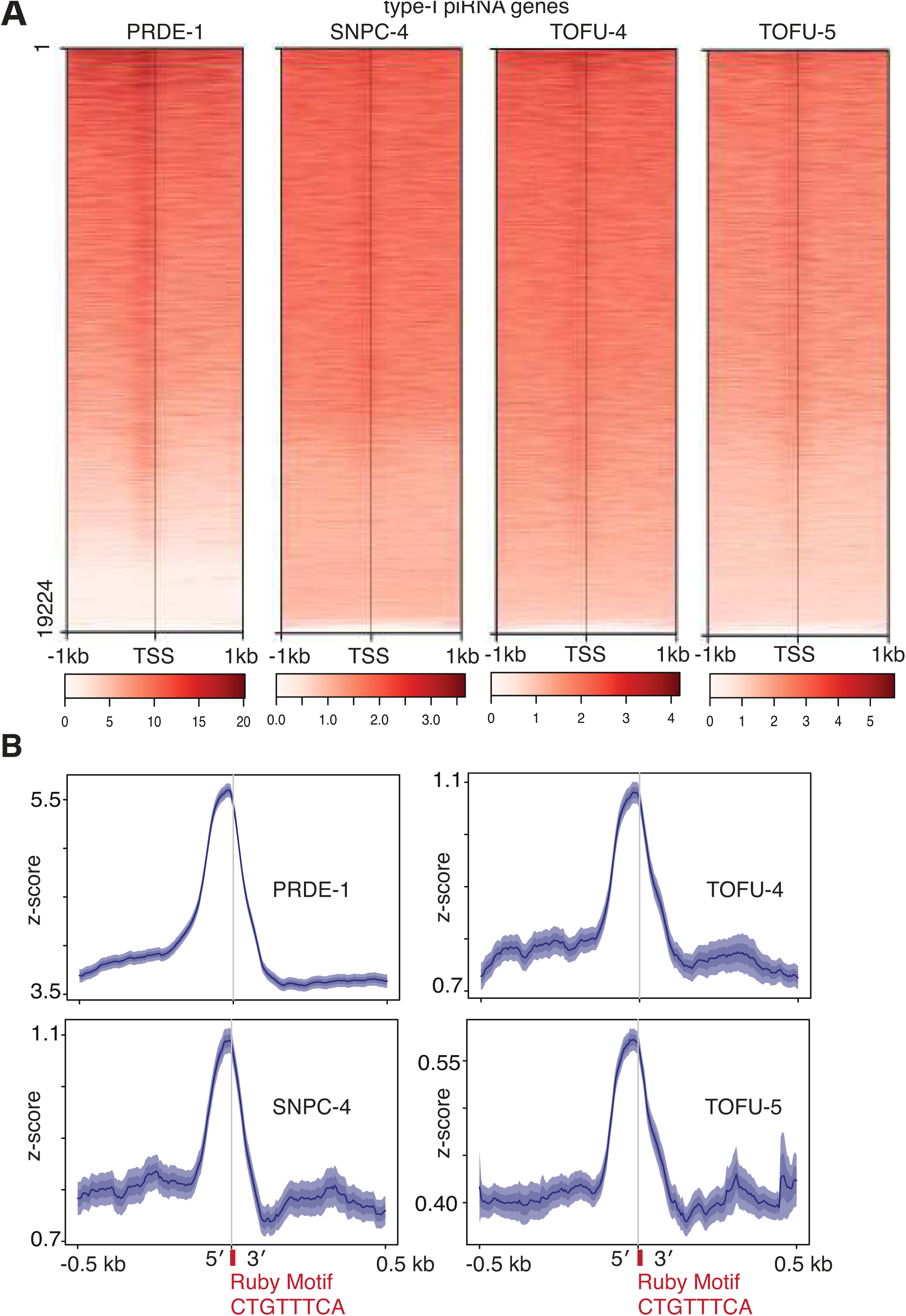
The USTC factors are enriched at the Ruby motif upstream of type I piRNA genes. (A) Heatmap of ChIP-seq binding profiles of USTC factors around type I piRNA transcription start sites. The BEADS normalisation score is plotted 1 kb upstream and downstream of the 1st U base of piRNAs. (B) Enrichment profile of USTC factors around the Ruby motif of type I piRNA genes within a 500 bps window.

### The binding of TOFU-5 to the piRNA clusters depends on the other USTC components

A previous study found that the concentration of SNPC-4 at 21U-RNA loci depends on PRDE-1 (Kasper et al. 2014). To investigate the genetic requirements of distinct USTC components for the binding of the 21U-RNA loci, we firstly examined whether the localisation of TOFU-5 to the piRNA loci depends on the presence of other USTC factors.

We crossed the TOFU-5∷GFP strain to *tofu-4(tm6157)* and *prde-1(mj207)* mutants, and found that TOFU-5 failed to form the sub-nuclear foci in germline cells in young adult animals (Fig. 4A). Additionally, ChIP-qPCR of TOFU-5 indicated that TOFU-5 does not bind to piRNA clusters in the absence of TOFU-4 or PRDE-1 (Fig. 4B). SNPC-4 is an essential gene required for the development of *C. elegans. snpc-4* mutant animals are embryonic or larval lethal (Kasper et al. 2014). We therefore crossed TOFU-5∷GFP animals with *snpc-4(tm4568/hT2)* balanced mutant animals and found that TOFU-5 fails to form the sub-nuclear foci in *snpc-4(tm4568)* homozygous mutant offspring of these animals (Fig. 4C). Remarkably, in *snpc-4(tm4568)* mutant, TOFU-5∷GFP accumulates in the germline syncytium, which was further confirmed by knocking down *snpc-4* by feeding RNAi (Fig. 4C). Both TOFU-5 and SNPC-4 contain a conserved SANT domain which may bind to DNA sequences (Boyer, Latek, and Peterson 2004). We constructed a TOFU-5(*SANT)∷GFP transgenic animal by deleting the SANT domain. Unlike wild-type TOFU-5, the TOFU-5(*SANT)∷GFP fails to form the piRNA foci, but is instead enriched in the germline syncytium (Fig. 4D). Consistently, TOFU-5(*SANT)∷GFP fails to bind to piRNA clusters, as shown by ChIP assay followed by real-time PCR (Fig. 4E). Proteins with SANT domains could heterodimerize (Horton et al. 2007). The similar cytoplasmic location of TOFU-5 (*SANT) and wild-type TOFU-5 in the absence of SNPC-4 suggests that the SANT domain of TOFU-5 might be required for TOFU-5’ interaction with SNPC-4. Together, we concluded that the ability of TOFU-5 to bind piRNA clusters and form the piRNA foci depends on other components of the USTC complex.

**Figure 4.**
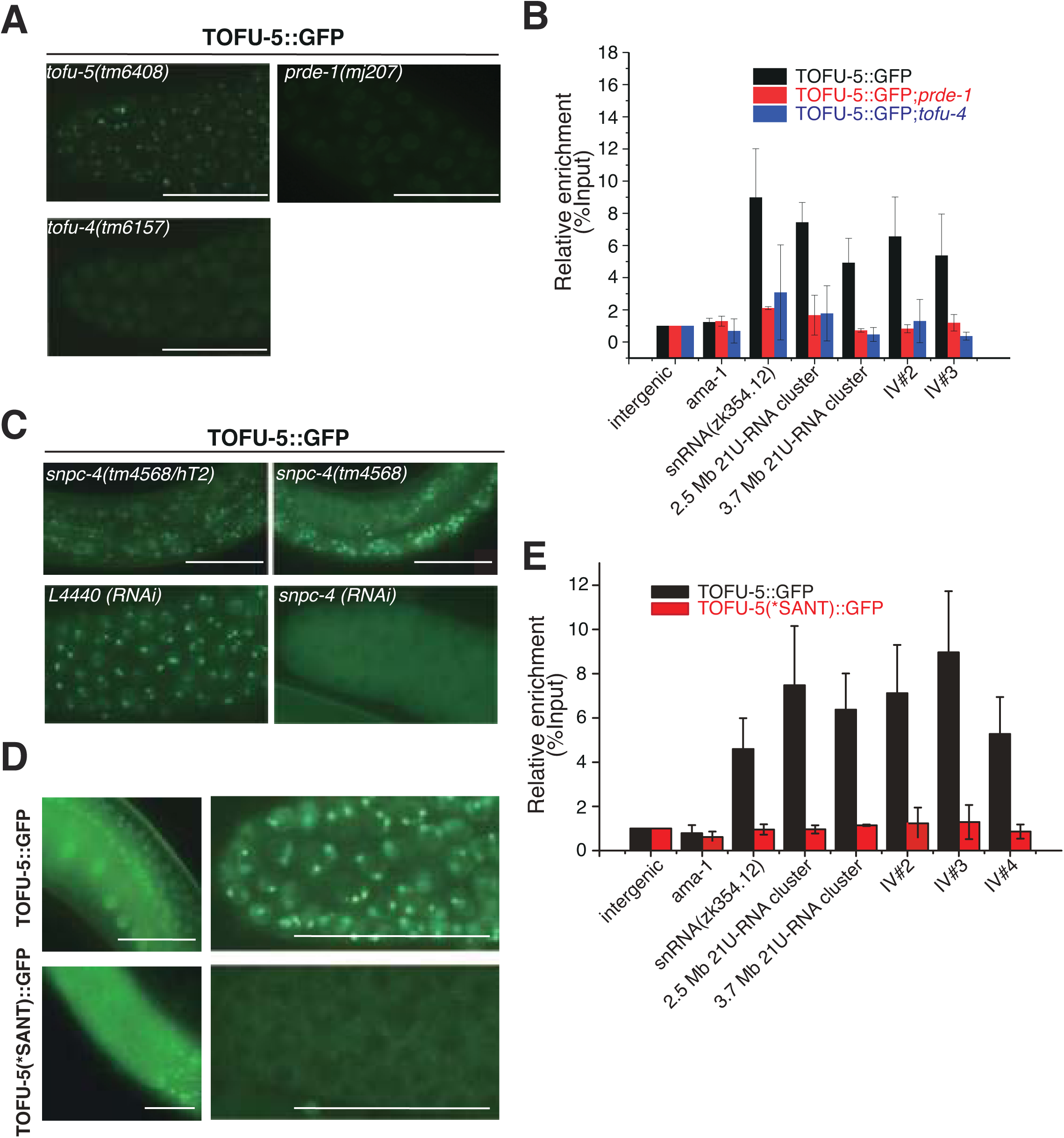
Genetic requirements of TOFU-5 binding to piRNA clusters. (A) Images of germline nuclei of animals expressing TOFU-5∷GFP. Scale bar, 20 μm. (B) Relative enrichment of TOFU-5 by ChIP assay with an anti-GFP antibody in indicated animals. n=3+/-1s.d. Images of germline nuclei of indicated animals. Primer sets are available in Supplementary Table S4. (C) TOFU-5∷GFP failed to localize to the nucleus in *snpc-4(-)* animals. Scale bar, 20 μm. (D) Images of germline nuclei of indicated animals. TOFU-5(*SANT)∷GFP failed to localize to the nucleus in distal germline. Scale bar, 20 μm. (E) Relative enrichment of TOFU-5∷GFP and TOFU-5(*SANT)∷GFP by ChIP assay with an anti-GFP antibody. (n=3+/-1s.d.)

### The binding of TOFU-4 to piRNA clusters depends on other USTC factors

Next, we examined whether the binding of TOFU-4 to the piRNA loci depends on the presence of other USTC components. We crossed TOFU-4∷GFP transgene into *prde-1(mj207)* mutants and found that PRDE-1 was also required for the formation of TOFU-4 piRNA foci (Fig. 5A) and the binding to piRNA clusters (Fig. 5B). We introduced TOFU-4∷GFP into *snpc-4(tm4568/hT2)* and *tofu-5(tm6408/hT2)* and found that TOFU-4 fails to form piRNA foci in *snpc-4(tm4586)* and *tofu-5(tm6408)* homozygous mutants, which was further confirmed by knocking down *snpc-4* by feeding RNAi (Fig. 5C). Therefore, we conclude that the binding of TOFU-4 to piRNA clusters depends on the presence of other USTC factors.

**Figure 5.**
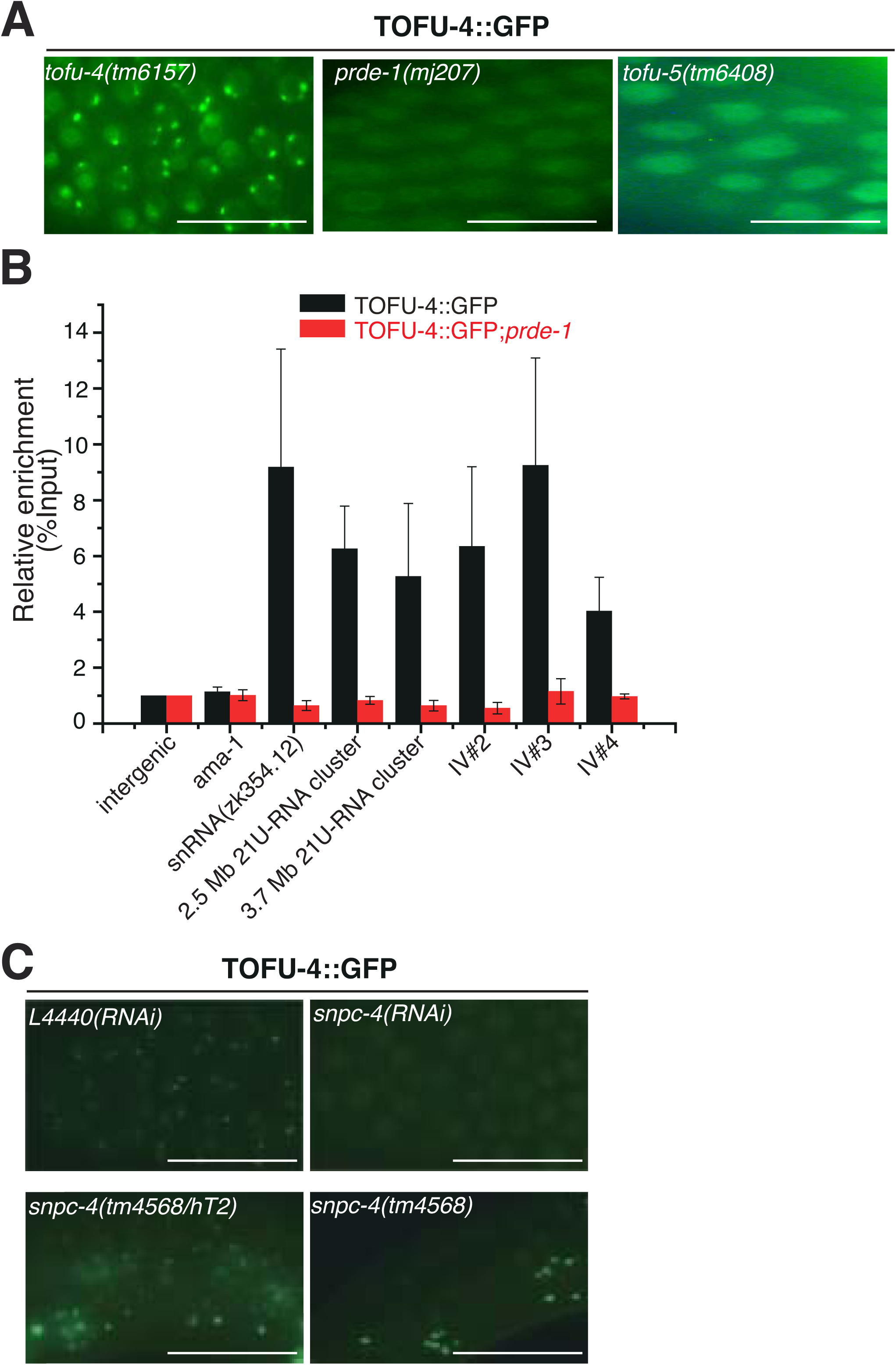
Genetic requirement of TOFU-4 binding to piRNA clusters. (A) Images of germline nuclei of indicated animals expressing TOFU-4∷GFP. Scale bar, 20 μm. (B) Relative enrichment of TOFU-4 by ChIp assay with an anti-GFP antibody. (n=3 +/-s.d.) Primer sets are available in Supplementary Table S4. (C) TOFU-4∷GFP fails to form nuclear foci in the germline upon RNAi feeding against SNPC-4.

### USTC binds to additional sets of non-coding genes

In addition to the piRNA clusters, SNPC-4 binds canonical SNAPc targets including RNA Pol II and RNA Pol III-transcribed non-coding RNA genes (Kasper et al. 2014). We therefore investigated whether PRDE-1, TOFU-4, TOFU-5, and SNPC-4 bound together also to other regions in the genome. Examining peaks for each factor outside of the piRNA clusters we found little evidence for PRDE-1 binding (Fig. 6). However, both SNPC-4 and TOFU-5 were bound to a number of small nuclear and small nucleolar RNA genes. As for TOFU-4 we did not observe many peaks but this might be a reflection of the signal to noise ratio of the TOFU-4 ChIP-seq experiment. Interestingly, TOFU-5 and SNPC-4 are also enriched on specific classes of transposable elements (Fig. S6). These results suggest that USTC might have acquired other functions in addition to promoting piRNA biogenesis (Fig. 7B).

**Figure 6.**
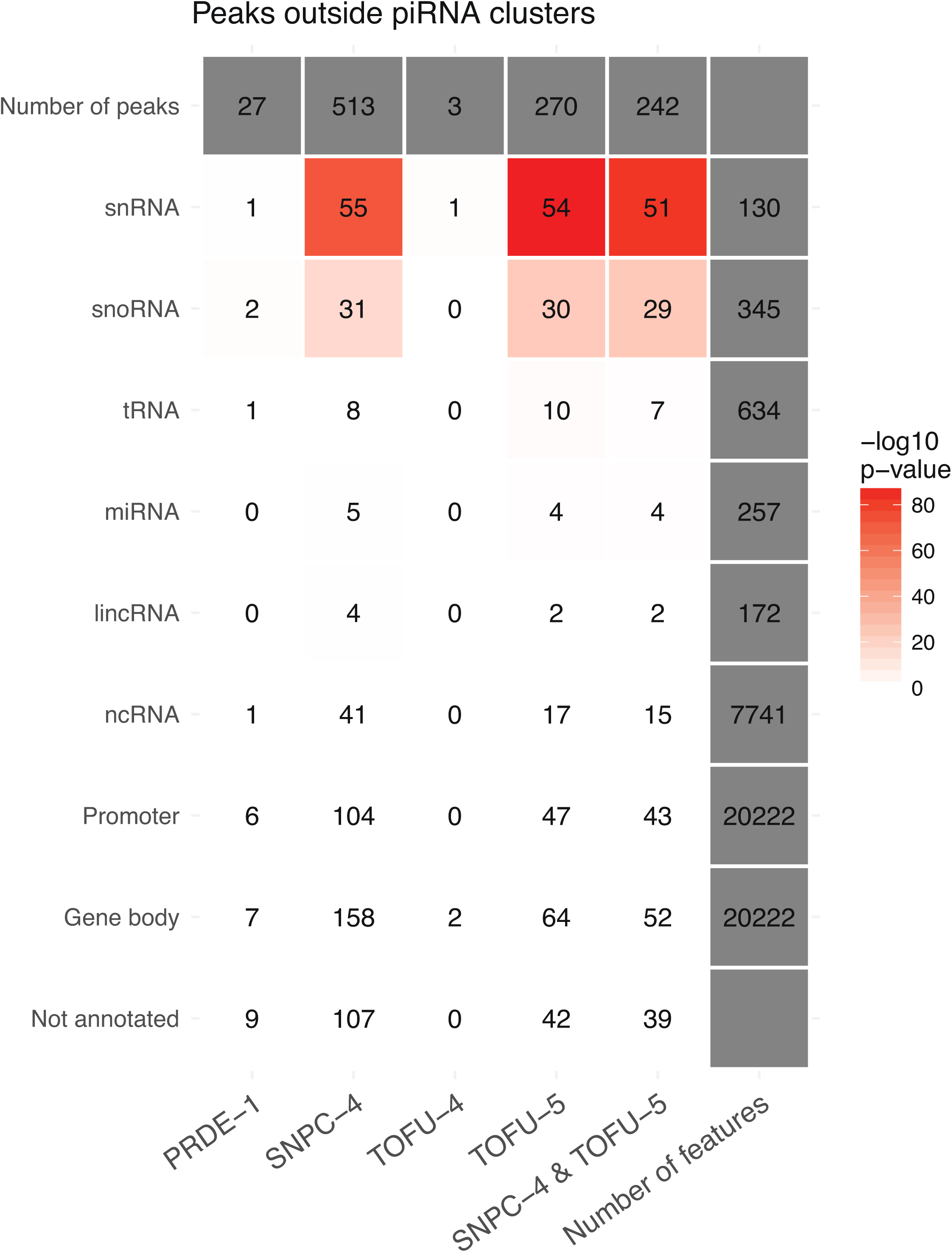
Binding sites of USTC factors outside of the piRNA clusters. Gene types of USTC binding sites are analyzed with selected annotations from Ensembl 92. The heatmap shows high confidence USTC factors binding sites overlaps with selected selected annotations from Ensembl 92. The bindings sites were detected using overlap between peak calls on individual replicates. The numbers denote direct overlaps between the peaks and annotated genomic loci, and colors identify the estimation of overlap significance (-log10 transformed p-value of hypergeometric test for overrepresentation).

**Figure 7.**
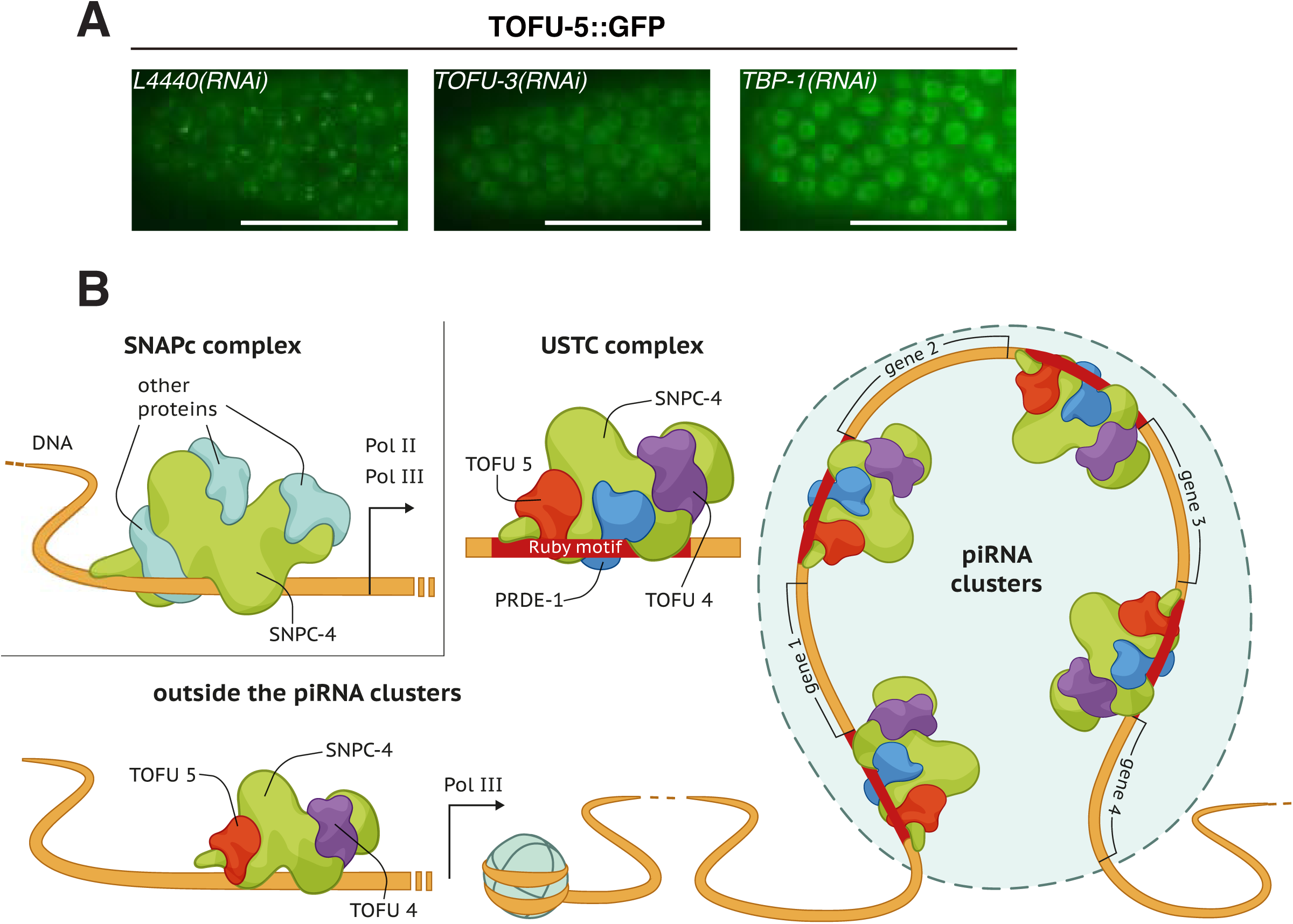
A model for the role of the USTC complex inside and outside of piRNA clusters. (A) Genetic reguirement for TOFU-5∷GFP in *tofu-3* and *tbp-1(*RNAi). (B) Proposed model for the mechanism of the USTC complex.

### Identification of TOFU-3 and TBP-1 as additional factors required for TOFU-5 binding to piRNA clusters

To further understand the function of the USTC complex in promoting piRNA transcription we searched for factors that are required for the formation of TOFU-5 sub-nuclear piRNA foci. We selected a number of candidate genes from our proteomic experiments and previous genome wide RNAi screens and carried out a focused candidate RNAi screen for TOFU-5 foci (Cecere et al. 2012; Goh et al. 2014) (Supplementary Table S2). Interestingly, we found that TOFU-3, a candidate from the genome wide RNAi screen, and TBP-1, a protein we identified through proteomics, are required for the formation of TOFU-5 piRNA foci (Fig. 7A; Fig. S7; Supplementary Table S2). However, we failed to identify any forkhead transcription factors in this screen, which previously had been shown to recognize the Ruby motif (Supplementary Table S2).

## DISCUSSION

Here, by a series of proteomics, imaging, and ChIP-seq experiments, we demonstrate that the four proteins PRDE-1, SNPC-4, TOFU-4, and TOFU-5 function as a complex and bind to the promoter sequences of individual piRNA transcription units. This complex localizes to sub-nuclear foci and exhibits concentrated binding across the two piRNA-rich domains on chromosome IV. We found that the two categories of piRNAs, classified by the presence of Ruby motif in their promoter region, may use different combination of the USTC factors for promoter recognition. While all of the USTC factors bound to the promoters of piRNA genes, TOFU-5 and SNPC-4 are also enriched on other classes of non-coding RNA (Fig. 7B).

### Using functional proteomic methods to identify the USTC complex

Previously, a number of forward genetic screens have been conducted and identified PRDE-1 and PID-1 as required for piRNA biogenesis in *C. elegans* (Weick et al. 2014; de Albuquerque et al. 2014). Using biochemical approaches, a Forkhead family transcription factor, *unc-130*, was shown to bind 21U-RNA promoters (Cecere et al. 2012). Additionally, a genome-wide RNAi screening identified seven TOFU genes that are engaged in distinct expression and processing steps of 21U-RNAs (Goh et al. 2014). However, since many of the genes are essential for the fertility or growth of *C. elegans*, these methods may not successfully recapitulate all the players and identify their distinct roles in piRNA biogenesis. Here, we combined a series of functional proteomic methods, including IP-MS and Y2H, and isolated a USTC complex that contain PRDE-1, SNPC-4, TOFU-4 and TOFU-5. These proteins can interact with each other and bind to the upstream promoter region of the piRNA transcription units. We further used chromatin immunoprecipitation and cell biology approaches and demonstrated a mutual dependency of the components of USTC complex in their ability of DNA binding and formation of the piRNA foci.

Interestingly, UNC-30 a member of the forkhead transcription factor family has been previously reported to bind piRNA gene promoters *in vitro* (Cecere et al. 2012). However, we did not observe UNC-30 or any other forkhead transcription factors’ enrichment with our mass spectrometry approaches. This might suggest alternative mechanisms that are potentially involved in piRNA biogenesis. In our study, we discovered a unique complex that differentially enriches over type I and type II piRNA genes and might engage the transcriptional machinery. Strikingly, in our USTC complex, we have characterized SNPC-4 protein which was previously reported in an ancient complex for ncRNA transcription with RNA pol II and pol III.

### The recognition of piRNA transcription units

*C. elegans* 21U-RNAs are classified into two types. Type I 21U-RNAs are predominantly transcribed from two broad regions from chromosome IV with an 8 nt upstream Ruby motif (CTGTTTCA) and a small YRNT motif. Type II 21U-RNAs lack Ruby motif or present outside of chromosome IV (Gu et al. 2012; Ruby et al. 2006). It has been previously shown that PRDE-1 and SNPC-4 are required for type I piRNA biogenesis (Weick et al. 2014). The forkhead transcription factors were reported to recognize the Ruby motif as well (Cecere et al. 2012). Here, we showed that, while the four factors of USTC complex bind type I piRNA promoters, TOFU-4 and PRDE-1 exhibit less binding activity to type II piRNA promoters. Whether this distinct pattern reflects Ruby-like sequences in type II piRNA promoters that exhibit weaker binding affinity or other protein factors are involved in the promoter recognition needs further investigation.

Remarkably, while SNPC-4 and TOFU-5 bind to overlapping sets of non-piRNA promoters, PRDE-1 lacked such binding sites. Therefore, the USTC complex may play a central role in the definition of piRNA transcription units and separate piRNA precursors from pre-mRNAs for downstream processing and maturation. RNA polymerase II transcribes protein-coding mRNA and also a variety of shorter noncoding RNAs, most notably spliceosomal U1 and U2 snRNAs (Lykke-Andersen and Jensen 2007; Egloff, O’, Reilly, and Murphy 2008). Although piRNA precursors and pre-mRNAs are both transcribed by RNA polymerase II, USTC may direct distinct transcription and processing machineries to piRNA units. It will be very interesting to examine the coupling between transcriptional regulation and processing and 3’ end trimming in the biogenesis of piRNAs. Experiments investigating the binding pattern of each component in the absence of other USTC factors will further enlighten our understanding of their mutual regulatory relationships (Lykke-Andersen and Jensen 2007; Egloff, O’Reilly, and Murphy 2008).

### TBP-1 and piRNA biogenesis

*tbp-1* encodes the *C. elegans* ortholog of the human TATA-box-binding protein (TBP), which plays important roles in transcriptional regulation. TBP-1 has been shown to provide TFIID-like basal transcription activity in human and *C. elegans* extracts, to bind specifically to a TATA box sequence, and to interact with TFIIA and TFIIB transcription factors. *tbp-1* activity is required for embryonic and larval development, as well as for normal rates of postembryonic growth. In *Drosophila*, TBP-1, also termed Moonshiner, drives the transcription of piRNA clusters (Andersen et al. 2017).

The function of *tbp-1* in piRNA biogenesis is yet unknown. Here, we showed that although TBP-1 did not accumulate in the piRNA foci (Fig. S7), it was required for the formation of the piRNA foci (Fig. 7A). We speculate that *tbp-1* and the Ruby motif may be required together for USTC binding to piRNA promoter. Here, TBP-1 may act as a bridge to bend the DNA so that the USTC complex comes closer to the piRNA promoter sequences. Further investigation on the roles of TBP-1 in piRNA biogenesis will shed light on the specific mechanistic understanding of the transcription regulation of piRNAs.

### Chromosome modification in piRNAs transcription

Both TOFU-5 and SNPC-4 contain SANT domains (Boyer, Latek, and Peterson 2004). SANT domains stand for ‘switching-defective protein 3 (Swi3), adaptor 2 (Ada2), nuclear receptor co-repressor (N-CoR), transcription factor (TF)IIIB. Sequence analysis of SANT domain indicates a strong similarity to the DNA-binding domain (DBD) of Myb-related proteins. SANT domains have been shown to couple histone-tail binding to enzymatic activity, including histone acetylation, de-acetylation, and ATP-dependent chromatin remodeling. Small deletions in the SANT domains may lead to a complete loss of function of the proteins. Here, we show that the deletion of the SANT domain of TOFU-5 relocalizes TOFU-5 from the nucleus to cytoplasm, disabled its binding ability to piRNA promoters, and alters the piRNA foci. The SANT domain-containing proteins can influence chromatin state, suggesting that SNPC-4 and TOFU-5 can modulate nucleosome organization and/or histone modifications in the 21U-RNA regions. Consistently, the 21U-RNA regions exhibited decreased nucleosome density in young adult animals (Cecere et al. 2012). It is unclear whether the presence of SANT domain is required for the cytoplasm-to-nucleus import of TOFU-5.

Chromatin modification plays important roles in small RNA biogenesis. In *Drosophila*, transcription of piRNA clusters-small RNA source loci is enforced through RNA polymerase II pre-initiation complex formation within repressive heterochromatin, in which the heterochromatin protein-1 variant Rhino recruits Moonshiner (Andersen et al. 2017). In *C. elegans*, two piRNA clusters on chromosome IV are nucleosome-depleted regions (Cecere et al. 2012). In our Y2H experiment, *set-6*, a gene encoding a putative H3K9 methyltransferase and *set-16*, a gene encoding a putative H3K4 methyltransferases were identified. However, these two proteins and other chromatin modification factors were unlikely required for the formation of piRNA foci (Supplementary Table S1). Therefore, we speculated that either the biogenesis of 21U-RNA is independent of chromatin modification processes or certain chromatin factors act together to promote the formation of piRNA foci. Further analysis of the properties of chromatin, including histone modifications, in the germline will be necessary to directly explore the relationship between the chromatin state, USTC complex, and 21U-RNA expression.

## Conclusion

Our work provides a foundation to understand piRNA transcription and biogenesis. Since the biological roles and regulation of *C. elegans* piRNAs (21U-RNAs) are still elusive, future characterization of the USTC complex is likely to shed new light on piRNA biology. Functional proteomics approaches will continue to be the key for future studies addressing the regulation of piRNA transcription and the coupling between the transcription and processing of piRNAs.

## Materials and Methods

### Strains

Bristol strain N2 was used as the standard wild-type strain. All strains were grown at 20° C unless specified. The strains used in this study are listed in Supplementary Table S3.

### Immunoprecipitation and mass spectrometry analysis

Anti-PRDE-1 antibody was produced at the Laboratory Animal Resources (EMBL Heidelberg). For this, two New Zealand White female rabbits were immunized with the recombinant full length PRDE-1 together with complete Freund’s adjuvant. After the priming immunization, four further booster immunizations were performed every 3 weeks. The harvested serum was purified by affinity purification using the recombinant full length PRDE-1.

Non-synchronized transgenic animals expressing TOFU-4∷GFP or TOFU-5∷GFP were resuspended in an equal volume of 2x lysis buffer (50 mM Tris-HCL pH 8.0, 300 mM NaCL, 10% glycerol, 1% Triton-X100, Roche^®^complete^TM^ EDTA-free Protease Inhibitor Cocktail, 10 mM NaF, 2 mM Na3V04) and lysed in a FastPrep-24 ^TM^ 5G Homogenizer. The supernatant of lysate was incubated with home-made anti-GFP beads for one hour at 4°C. The beads were then washed three times with cold lysis buffer. GFP immuno-precipitates were eluted by chilled elution buffer (100 mM Glycine-HCI pH 2.5). About 1/8 of the eluates were subjected to the western blotting analysis. The rest eluates were precipitated in TCA or cold acetone and dissolved in 100 mM Tris, pH 8.5, 8 M urea. The proteins were reduced with TCEP, alkylated with IAA, and finally digested with trypsin at 37°C overnight. The LCMS/MS analysis of the resulting peptides and the MS data processing approaches were conducted as previously described (G. Feng et al. 2017). Briefly, samples were digested with trypsin and analysed on an LTQ Orbitrap Velos Pro (Thermo Fisher Scientific) LC/MS-MS system. Data were searched with the Mascot (v 2.2.03, Matrix Science) against the Uniprot *C.elegans* database and unique peptides were analysed with Scaffold 3.

### Construction of plasmids and transgenic strains

For *TOFU-4∷GFP, a* TOFU-4 promoter and CDS region was PCR amplified with the primers <u>5’-GCAGA TCTTC GAATG CATCG AATTG AAAAT GAGAA AAAAC TG-3’</u> and <u>5’-ATAGC TCCAC CTCCA CCTCC GAATT CTGCG TCTAC CATCA-3’</u> from N2 genomic DNA. A GFP∷3xFLAG region was PCR amplified with the primers <u>5’-GGAGG TGGAG GTGGA GCTAT GAGTA AAGGA GAAGAA C-3’</u> and <u>5’-TCACT TGTCA TCGTC ATCCT-3’</u> from plasmid pSG085. A TOFU-4 3’ UTR was PCR amplified with the primers <u>5’-AGGAT GACGA TGACA AGTGA AAGAA TTTTT ATCCG AGTCA C-3’</u> and <u>5’-AGATA TCCTG CAGGA ATTCC GAAAC TTGAT TTTCA AAATT TA-3’</u> from N2 genomic DNA. The ClonExpress^®^ MultiS One Step Cloning Kit (Vazyme C113-02, Nanjing) was used to insert *TOFU-4∷GFP∷3xFLAG* fusion gene into pCFJ151 vector. The transgene was integrated onto the C. chromosome II by MosSCI method (Frøkjaer-Jensen et al. 2008).

For *TOFU-5∷GFP, a* TOFU-5 promoter and CDS region was PCR amplified with the primers <u>5’-GCAGA TCTTC GAATG CATCG TTGGT TCAAC CTGGA ATTAA-3’</u> and <u>5’-ATAGC TCCAC CTCCA CCTCC ATTGG ACGAT GTAGA TGGCT G-3’</u> from N2 genomic DNA. A GFP∷3xFLAG region was PCR amplified with the primers <u>5’-GGAGG TGGAG GTGGA GCTAT GAGTA AAGGA GAAGA AC-3’</u> and <u>5’-AAAAA TTTCA CTTGT CATCG TCATC CTTGT-3’</u> from plasmid pSG085. A TOFU-5 3’-UTR was PCR amplified with the primers <u>5’-CGATG ACAAG TGAAA TTTTT ATTAA TTTTT</u> and <u>5’-AGA TATCC TGCAG GAATT CCCAT GGCTT GGAAT TGAAG-3’</u> from N2 genomic DNA. The ClonExpress^®^ MultiS One Step Cloning Kit (Vazyme C113-02, Nanjing) was used to insert *TOFU-5∷GFP∷3xFLAG* fusion gene into pCFJ151 vector. The transgene was integrated onto the *C. elegans* chromosome II by MosSCI system.

For *TOFU-5(*SANT)∷GFP*, plasmid was PCR amplified with the primers <u>5’-TCGCA GCTTC ATCGA AGAGA TAAAG TAAGT-3’</u> and <u>5’-TCTCT TCGAT GAAGC TGCGA CCTTC TGCGA-3’</u> from *TOFU-5∷GFP∷3xFLAG* plasmid. The transgene was integrated onto the *C. elegans* chromosome II by MosSCI system.

### RNAi

RNAi experiments were conducted as previously described (Timmons, Court, and Fire 2001). Images were collected using a Leica DM2500 microscope.

### ChIP

Chromatin immunoprecipitation (ChIP) experiments were performed as previously described with hypochlorite-isolated embryos or young adults (Guang et al. 2010). Animals were crosslinked in 2% formaldehyde for 30 min. Fixation was quenched with 0.125 M glycine for 5 min at room temperature. After crosslinking, samples were resuspended in FA buffer (50 mMTris/HCl [pH 7.5], 1 mM EDTA, 1% Triton X-100, 0.1% sodium deoxycholate, and 150 mM NaCl) with proteinase inhibitor tablet (Roche no. 04693116001) and sonicated for 20 cycles at medium output (each cycle: 30s ON and 30s OFF) with a Bioruptor 200. Lysates were precleared and then immunoprecipitated with 1.5 μl anti-GFP anti-body (Abcam, ab290) for TOFU-4, TOFU-5, and SNPC-4, 5 μl anti-PRDE-1 for PRDE-1 overnight at 4°C. Antibody-bound complexes were recovered with Dynabeads^®^ Protein A (Thermo, 10002D). Following to extensive squential washes with 150, 500 and 1M NaCl, DNA was treated with RNAse (Roche) and ProK (New England Biolabs). Finally resulting DNA samples were purified with QIAquick^®^ PCR Purification Kit (QIAGEN, 28104).

### ChIP-seq

The DNA samples from ChIP experiment were sent to in-house sequencing for library preparation and sequencing. Briefly, 10-300 ng ChIP DNA was combined with End Repair Mix and incubated at 20°C for 30 min, followed by purification with a QIAquick PCR Purification Kit (Qiagen). The DNA were then incubated with A-Tailing Mix at 37°C for 30 min. The 3’ ends adenylated DNA was incubated with adapter in the ligation mix at 20°C for 15 min. The adapter-ligated DNA was amplified by several rounds of PCR amplification and purified by a 2% agarose gel to recover the target fragments. The average molecule length was assayed by using the Agilent 2100 bioanalyzer instrument (Agilent DNA 1000 Reagents). The library was quantified by quantitative real-time PCR (QPCR) (TaqMan Probe). The libraries were further amplified on cBot to generate the clusters on the flowcell and sequenced with a single end 50 method on a HiSeq1500 System.

### Alignment to reference genome for ChIP-seq data

Chip-seq libraries were sequenced using lllumina HiSeq. Reads were aligned to the cel 1 assembly of the *C. elegans* genome using BWA v. 0.7.7 (Li and Durbin, 2010) with default settings (BWA-backtrack algorithm). The SAMtools v. 0.1.19 ‘view’ utility was used to convert the alignments to BAM format. Normalized ChIP-seq coverage tracks were generated using the BEADS algorithm (Cheung et al 2011).

### Summed ChIP-seq Input and in-house blacklist

We generated summed input BAM files by combining good quality ChIP-seq input experiments from different extracts (8 experiments for formaldehyde and 5 experiments for EGS extracts). The same summed inputs were used for BEADS normalisation and peak calls. We observed that despite using input files for MACS2 (Zhang et al. 2008) and filtering against modENCODE, some regions of high signal in input were still called as peaks. To overcome this problem, we created an in-house blacklist by running MACS2 with default settings and no input mode. The blacklist regions were refined by discarding region with MACS2 score below 100 and clustering peaks within 500bp. This procedure created 90 new regions in addition to 122 already covered by modENCODE blacklist.

### Peak calls

ChIP-seq peaks were called using MACS2 v. 2.1.1 (Feng and Liu, 2011) with permissive 0.05 q-value cut-off and fragment size of 150bp against summed ChIP-seq input. To generate sharp peak calls set, we obtained peaks summits and extended them 150bp upstream and downstream, creating 300pb regions around summit calls. Further, we combined ChIP-seq replicates by intersecting these regions and setting the final peaks size back to 300bp. Finally, peaks overlapping non-mappable (GEM-mappability < 25%) or blacklisted regions were discarded. This produced sharp, uniform peaks suitable for further quantitative analyses.

### Peak calls annotation

PRDE-1, SNPC-4, TOFU-4 and TOFU-5 peak calls were classified using selected annotations from Ensembl v92. The peak is assigned to given class if it directly overlaps with annotated loci. The assignments are exclusive (a peak can be assigned only to a single class), giving following order of priority: snRNA, snoRNA, tRNA, miRNA, lincRNA, ncRNA, promoter, gene body. Promoters are defined as 500 bases upstream of annotated transcription start site (TSS). The overlap significance was estimated using hypergeometric test for overrepresentation.

### ChIP-seq data aggregation and visualization

SeqPlots (Stempor & Ahringer 2016) software was used to visualise PRDE-1, SNPC-4, TOFU-4 and TOFU-5 ChIP-seq profiles over Ruby motif loci, piRNAs precursors, snRNAs and tRNAs average as average aggregated plots and heatmaps. The IGV Genome Browser (Robinson et al. 2011) was applied to visualise signal genome-wide and on piRNA clusters.

### ChIP-seq signal quantifications

To quantify differences in PRDE-1, SNPC-4, TOFU-4 and TOFU-5 binding between piRNA clusters and somatic chromosomes we quantified BEADS normalised, log2 scaled signal in 1 KB bins divided into piRNA clusters and chromosomes I, II, III & V. The signal was obtained using bigWigSummary utility from Kent library (Kent et al. 2010) implemented in rtracklayer package in R. Then, signal was represented as overlaid violin plot (showing signal distribution) and Tukey box plot (showing estimation of statistical significance of difference between medians as notches) (Tukey, 1977).

### Statistics

Bar graphs with error bars were presented with mean and standard deviation (s.d.). All the experiments were conducted with independent *C. elegans* animals for indicated N times. Statistical analysis was performed with two-tailed Student’s t-test.

### Data availability

All raw and normalized seguencing data has been deposited to GEO under the submission name GSE112682.

### piRNA Gene Annotations

piRNA annotations were downloaded from the piRBase online database (http://www.regulatoryrna.org/database/piRNA/). Genomic coordinates of piRNA genes were obtained by Samtools against *C. elegans* ce10 genome assembly. Type II piRNA genes were obtained from a previous publication (Gu et al. 2012). Type I piRNA gene lists were created by filtering the piRBase annotations with type II piRNA genes.

## Acknowledgments

We are grateful to the members of the Guang lab and Miska lab for their comments. Especially, we are grateful to Lisa Lampersberger and Dr. Giulia Furlan for the critical reading of the manuscript. We thank the Laboratory Animal Resources, the Protein Expression and Purification and the Proteomics Core Facilities at EMBL Neidelberg for materials and technical support. We are grateful to the International *C. elegans* Gene Knockout Consortium, and the National Bioresource Project for providing the strains. Some strains were provided by the CGC, which is funded by NLN Office of Research Infratructure Programs (P40 OD010440). This work was supported by grants from the Chinese Ministry of Science and Technology (2017YFA0102903 and 2014CB84980001), the National Natural Science Foundation of China (Nos. 81501329, 31371323, 31671346, and 91640110), Anhui Natural Science Foundation (No. 1608085MC50), Beijing Municipal Science and Technology Commission (fund for cultivation and development of innovation base), Cancer Research UK (C13474/A18583, C6946/A14492) and the Wellcome Trust (104640/Z/14/Z, 092096/Z/10/Z). A.C.B. is supported by a Marie Skladowska Curie Post-Doctoral Fellowship.

## Author contributions

C.W., S.G., A.C.B, J.A. and E.A.M. wrote the manuscript with input from the co-authors. A.C.B., C.W. and P.S. analyzed the sequencing data. C.W., A.K., A.C.B., X.F., H.M., C.Z., W.L., Y.Y., and M.D. performed the experiments. O.B. and C.Z. generated the antibodies and performed the MS run and analysis. E.A.M. and S.G. conceived the project and coordinated the studies.

## SUPPLEMENTARY FIGURE LEGENDS

**Figure S1.**
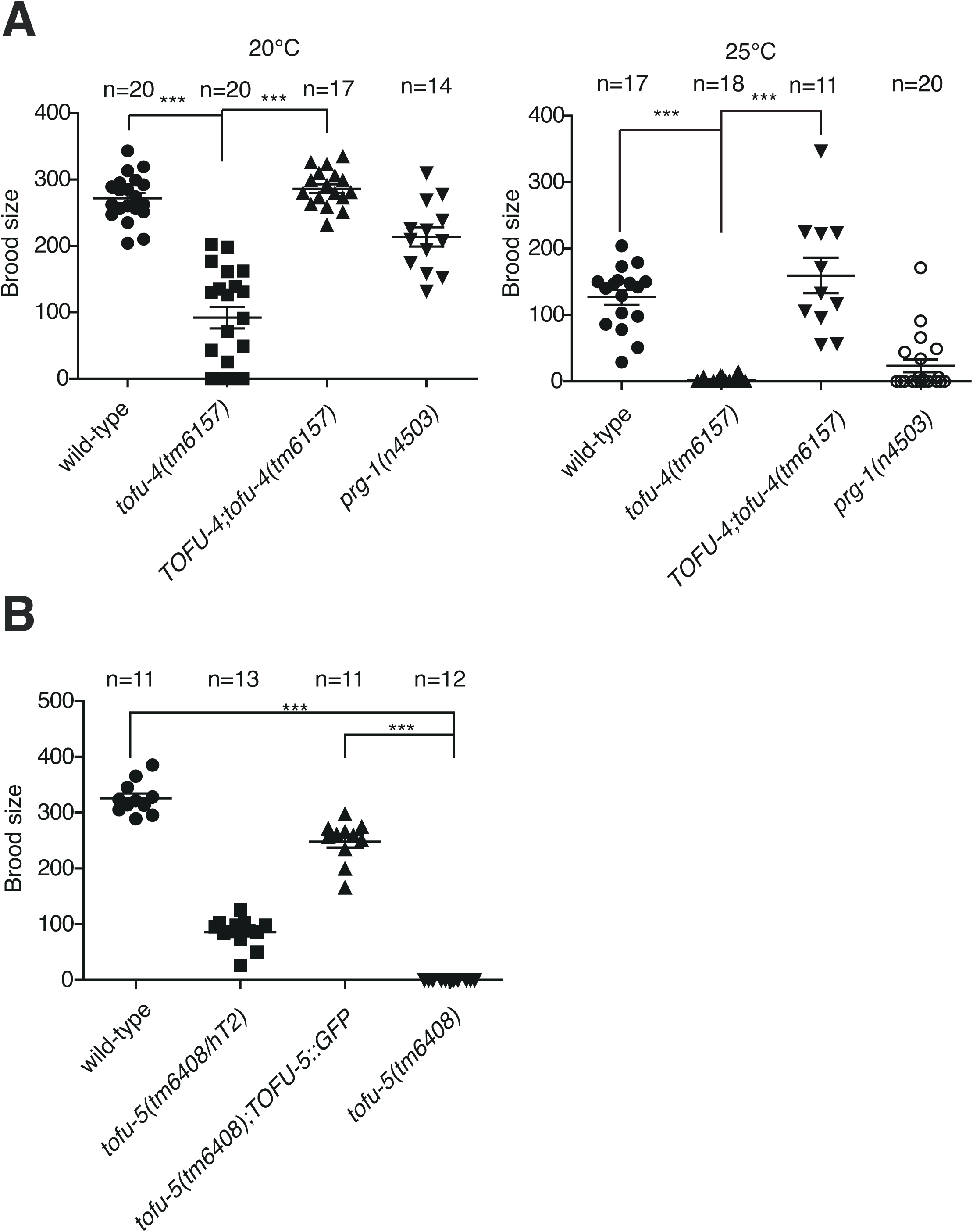
TOFU-4∷GFP and TOFU-5∷GFP are functional. (*A*) TOFU-4 is reguired for normal fertility, TOFU-4∷GFP rescued the progeny defect of *tofu-4(tm6157)* mutant. Average brood size of wild-type, *tofu-4(tm6157), prg-1(n4503)* mutant stains and *tofu-4* rescue lines at 20°c and 25°C. (*B*) TOFU-5 is required for normal fertility, TOFU-5∷GFP rescued the sterile phenotype of *tofu-5(tm6408)* mutant. Average brood size of wild-type and *tofu-5(tm6408)/hT2, TOFU-5-(tm6480)* rescue line, *tofu-5(tm6408)* at 20°C. (n) Number of parental adults used.

**Figure S2.**
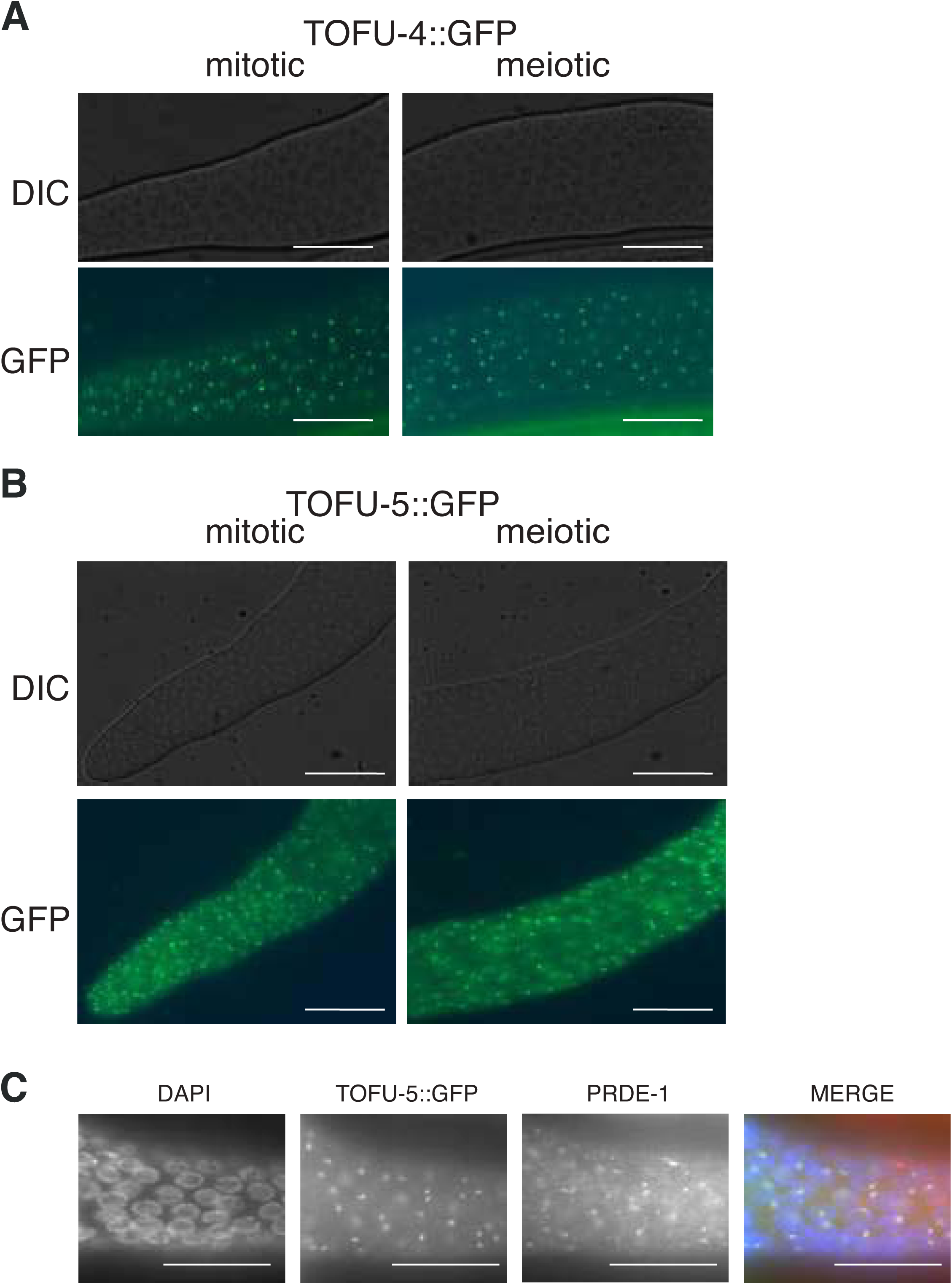
Expression pattern of TOFU-4, TOFU-5, and PRDE-1. (*A*) Image of TOFU-4 in cut gonad. (*B*) Image of TOFU-5 in cut gonad. (*C*) Colocalization of TOFU-5 and PRDE-1. Scale bar, 20 μm.

**Figure S3.**
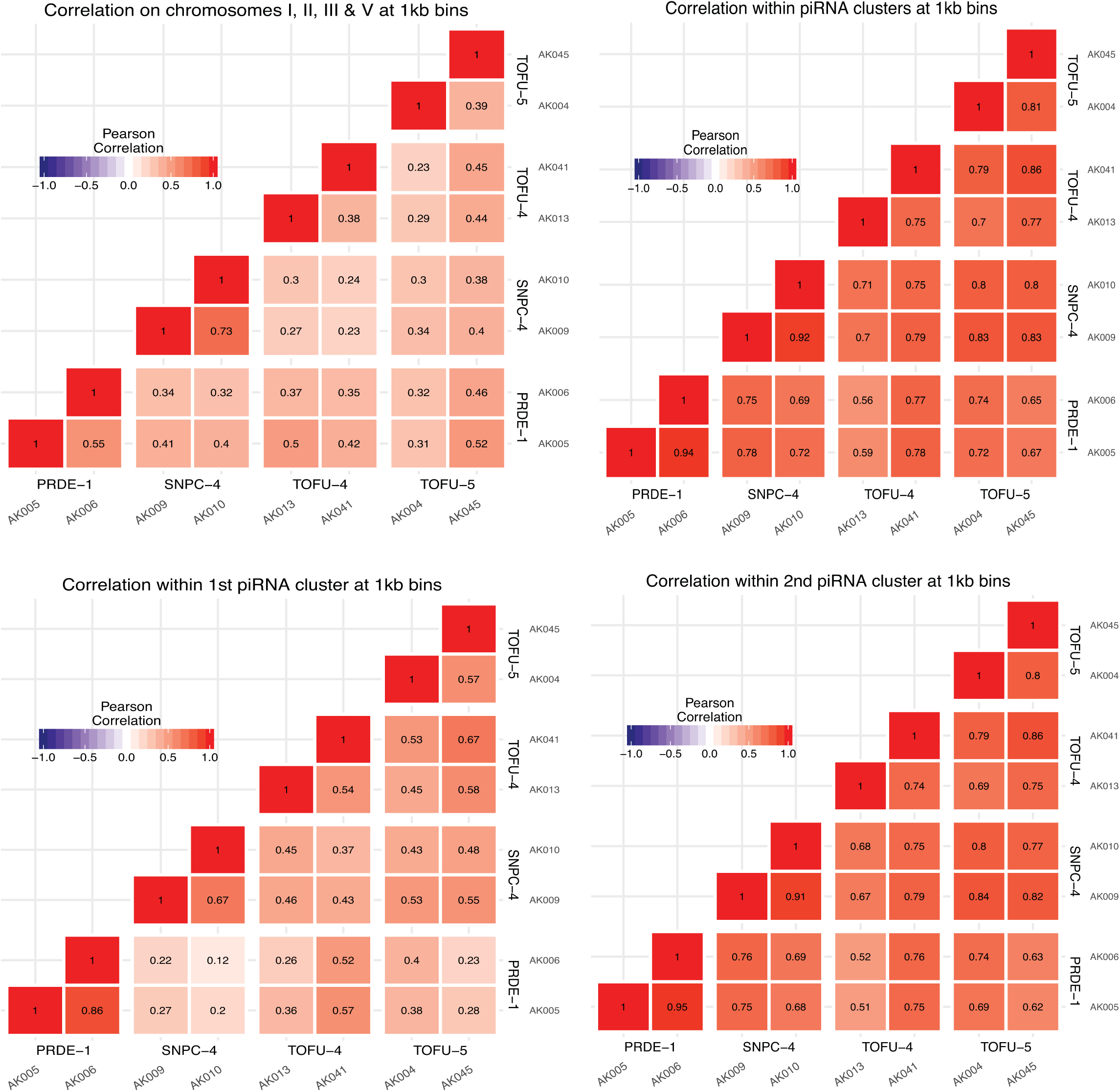
Pearson Correlation between ChIP-seq replicates. Pearson correlation between the USTC ChIP-seq replicates using 1 kb bins.

**Figure S4.**
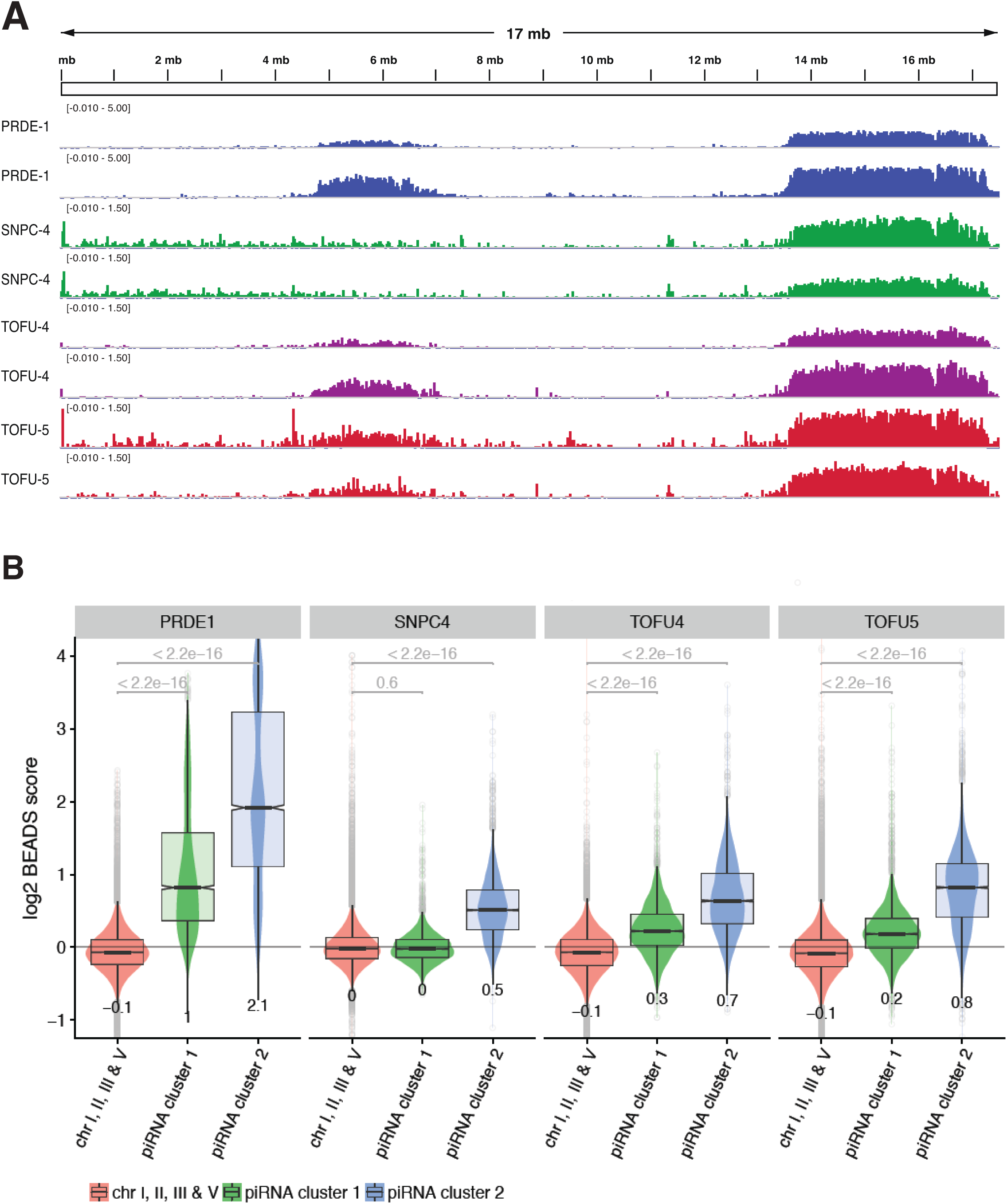
The USTC complex binds piRNA clusters on chromosome IV. (*A*) log2 fold enrichment of ChIP-seq individual replicates from the USTC complex on the piRNA clusters. (*B*) Log2 BEADS scores of the USTC complex in piRNA cluster I and II of chromosome IV over the genome average.

**Figure S5.**
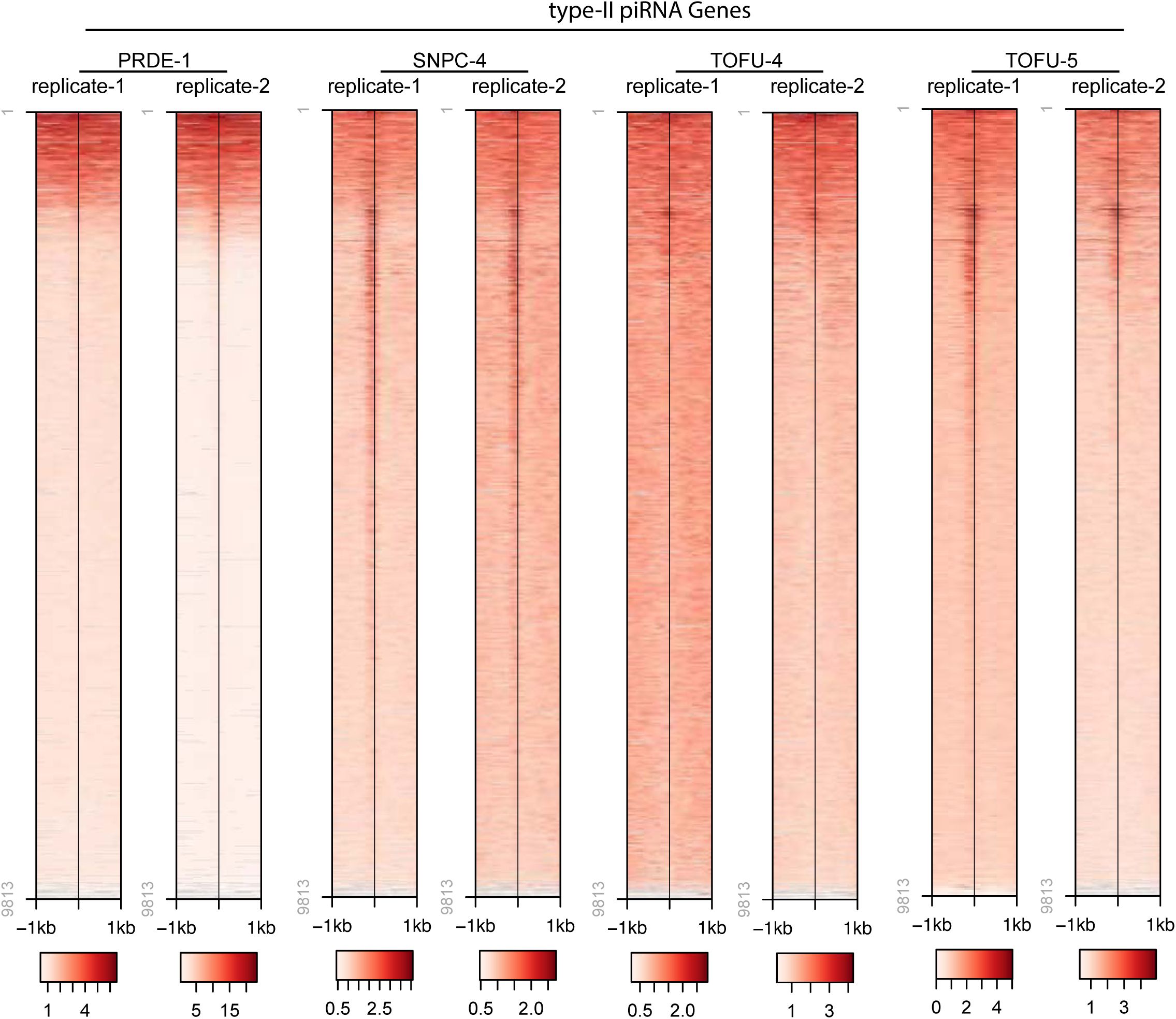
Heatmap profiles of the USTC complex on type II piRNA genes. Fold enrichment of the USTC complex ChIP-seq over type II piRNA genes. The BEADS score was plotted 1 kb upstream and downstream of the 1st U base of piRNAs.

**Figure S6.**
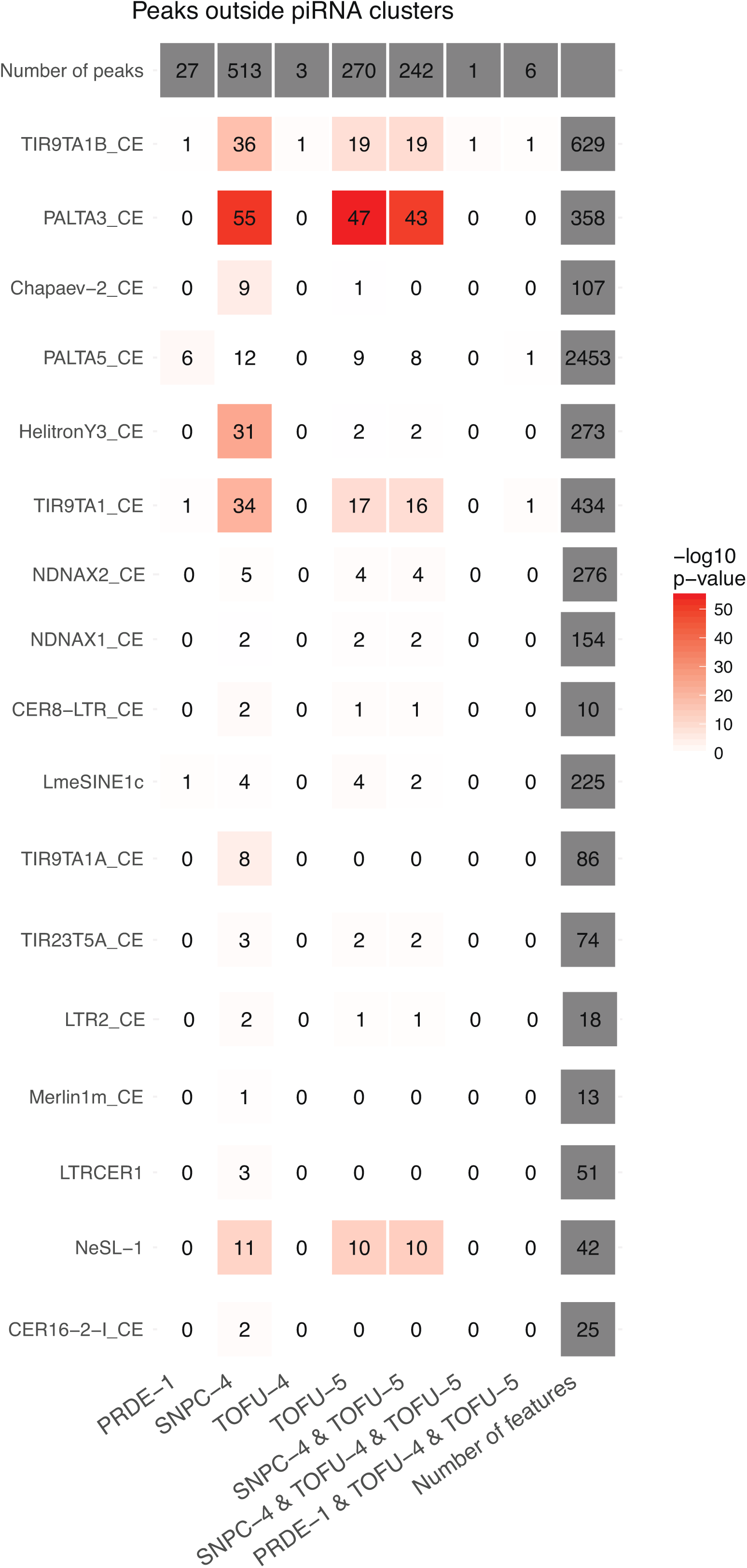
USTC factor association with repeat elements. Dfam2.0 repeat annotations were used for the analysis. Elements that showed at least one intersection with the USTC factors are shown

**Figure S7.**
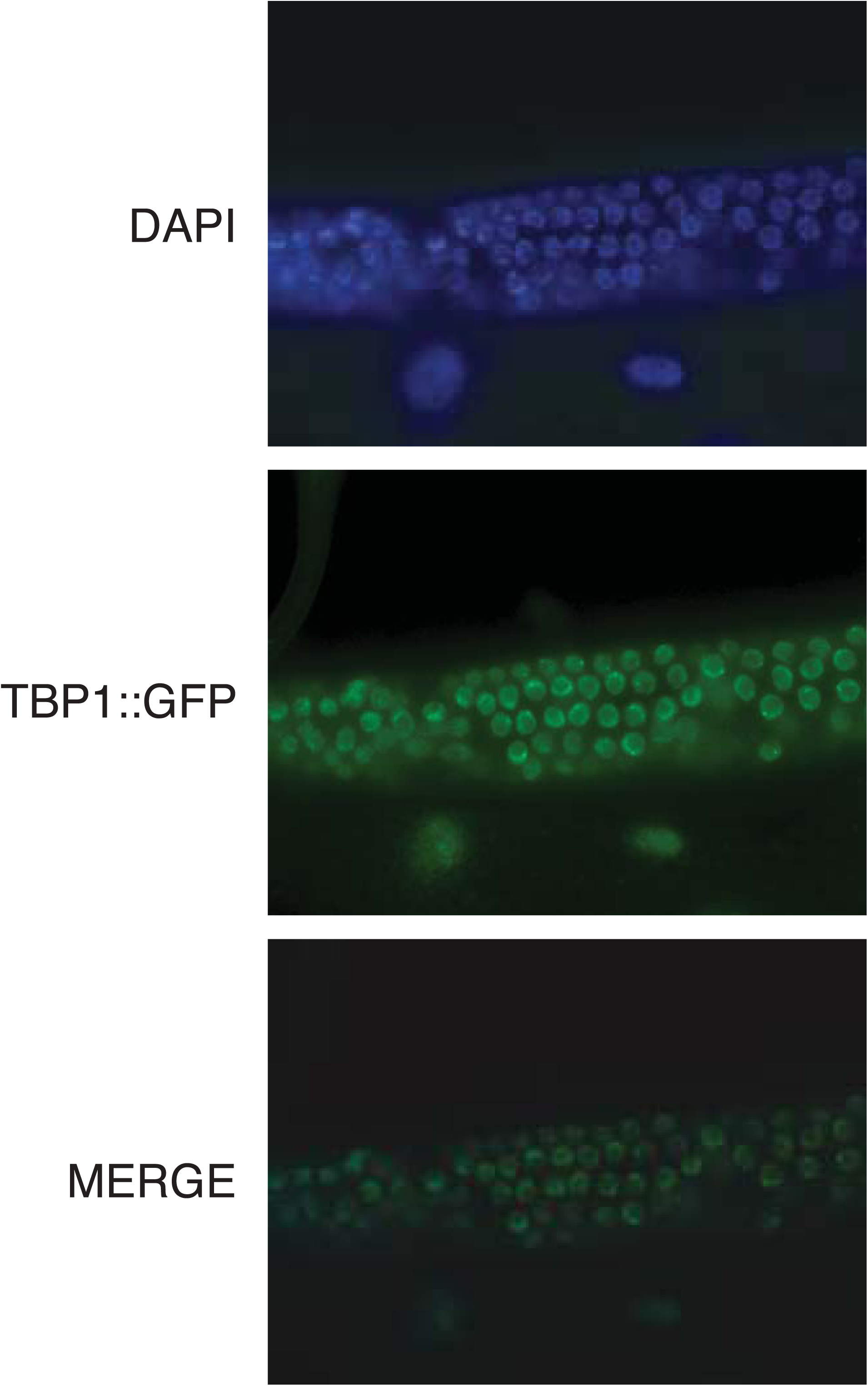
Expression pattern of TBP-1. Image of TBP-1 ∷GFP in the dissected gonad. Scale bar, 20 μm.

